# An inducible CRISPR-ON system for controllable gene activation in human pluripotent stem cells

**DOI:** 10.1101/106112

**Authors:** Jianying Guo, Dacheng Ma, Rujin Huang, Jia Ming, Min Ye, Kehkooi Kee, Zhen Xie, Jie Na

## Abstract

Human pluripotent stem cells (hPSCs) are an important system to study early human development, model human diseases, and develop cell replacement therapies. However, genetic manipulation of hPSCs is challenging and a method to simultaneously activate multiple genomic sites in a controllable manner is sorely needed. Here, we constructed a CRISPR-ON system to efficiently upregulate endogenous genes in hPSCs. A doxycycline (Dox) inducible dCas9-VP64-p65-Rta (dCas9-VPR) transcription activator and a reverse Tet transactivator (rtTA) expression cassette were knocked into the two alleles of the *AAVS1* locus to generate an iVPR hESC line. We showed that the dCas9-VPR level could be precisely and reversibly controlled by addition and withdrawal of Dox. Upon transfection of multiplexed gRNA plasmid targeting the *NANOG* promoter and Dox induction, we were able to control *NANOG* gene expression from its endogenous locus. Interestingly, an elevated *NANOG* level did not only promote naïve pluripotent gene expression but also enhanced cell survival and clonogenicity, and it enabled integration of hESCs with the inner cell mass (ICM) of mouse blastocysts *in vitro*. Thus, iVPR cells provide a convenient platform for gene function studies as well as high-throughput screens in hPSCs.

## Introduction

Human pluripotent stem cells (hPSCs), including human embryonic stem cells (hESCs) and human induced pluripotent stem cells (hiPSCs), are capable of self-renewal indefinitely and have the potential to differentiate into all cell types in the human body. Therefore this system offers a useful platform to study early human embryogenesis and a potential cell source for regenerative medicine. Moreover, functional cells derived from hESCs can be used to model human diseases in the context of drug toxicity tests and new drug development. These applications rely on methods to precisely control gene expression. However, because of difficulties in culture and transfection, targeted regulation of gene expression in hPSCs remains a technically challenging task. A method for efficient, rapid, and controllable gene activation is sorely needed.

Recently, the clustered regularly interspaced short palindromic repeat (CRISPR)/Cas9 system emerged as a powerful and versatile tool for genome editing (Wiedenheft et al. 2012). CRISPR was initially discovered as the adaptive immune system of bacteria and archaea (Wiedenheft et al. 2012). In response to viral and plasmid infection, bacteria and archaea could cut and degrade the foreign DNA recognized by a matching spacer RNA with the help of the Cas9 enzyme (Wiedenheft et al. 2012). CRISPR was rapidly transformed to a genome editing tool, and it has been shown to work in a wide range of systems, from plants to human cells, since the Cas9 nuclease can be directed easily to virtually anywhere in the genome using a short guide RNA and cutting the target DNA (Hsu et al. 2014). In pluripotent stem cells, the CRISPR system has been used to perform highly efficient gene knock-out and knock-in studies (Hsu et al. 2014). In addition to genome editing, a nuclease inactivated Cas9 (dCas9) was developed (Gilbert et al. 2014). By fusing dCas9 with transcription activators and repressors, such as VP64, and KRAB (Balboa et al. 2015; Gilbert et al. 2014; Mandegar et al. 2016; Genga et al. 2015), or with epigenetic modifiers, such as the catalytic domain of acetyltransferase p300 (Hilton et al. 2015) and Tet (ten eleven translocation) dioxygenase (Xu et al. 2016), one can use the CRISPR system to activate or inhibit gene expression or modify the histone and DNA methylation status at the desired locus.

Because of its potential applications in regenerative medicine, random insertion of foreign DNA into the genome of hPSCs should be avoided, since this may cause harmful mutations. The Adeno-Associated Virus Integration Site 1 (*AAVS1*) locus resides in the first intron of the *PPP1R12C* gene and has been used as a safe harbor for transgene integration (Smith et al. 2008; Hockemeyer et al. 2009; Lombardo et al. 2011; Qian et al. 2014; Zhu et al. 2014; Genga et al. 2015). Here we generated an iVPR hESC line by knocking-in the inducible dCas9-VPR system into the two alleles of the *AAVS1* locus. Detailed characterization of the iVPR hESC demonstrated that dCas9-VPR protein could be induced by Dox within 12 hours and disappear after Dox withdrawal. An inducible *NANOG* overexpression line (iNANOG) was established based on the iVPR system. We found a significant increase in NANOG protein after Dox induction. INANOG cells upregulated naïve pluripotency genes and were able to grow for a significant length of time in a naïve state medium containing ERK and GSK3 inhibitors and human LIF. The iVPR system can be a valuable system to control gene expression from endogenous loci and serve as platform for genome wide screens to identify new genes that can regulate stem cell self-renewal and differentiation.

## Results

### DCas9-VPR mediated robust ectopic and endogenous gene activation in human cell lines

To construct a robust and tunable gene activation system in hPSCs, we first compared the activation efficiency of dCas9-VPR (Chavez et al. 2015) with dCas9-VP64 (Kearns et al. 2014) and the Doxycycline (Dox) inducible Tet-On transactivator (rtTA) (Fig. 1A). We constructed plasmids to express gRNA targeting the TetO sequence (gTetO), and tested the ability of dCas9-VPR+gTetO or dCas9-VP64+gTetO to activate the synthetic TRE promoter driving enhanced blue fluorescent protein expression (TRE-BFP) in 293FT cells (Fig. 1A). The Tet transactivator (rtTA) was used as positive control (Fig. 1B). DCas9-VPR strongly activated BFP fluorescence, 43.1% of cells were BFP positive, while in the rtTA+Dox and dCas9-VP64 groups, only 28.2% and 5.8% of cells activated BFP, respectively (Fig. 1C and D). Moreover, dCas9-VPR resulted in the strongest mean BFP fluorescence intensity, indicating that it is the strongest activator among the three (Fig. 1D).

**Figure 1.**
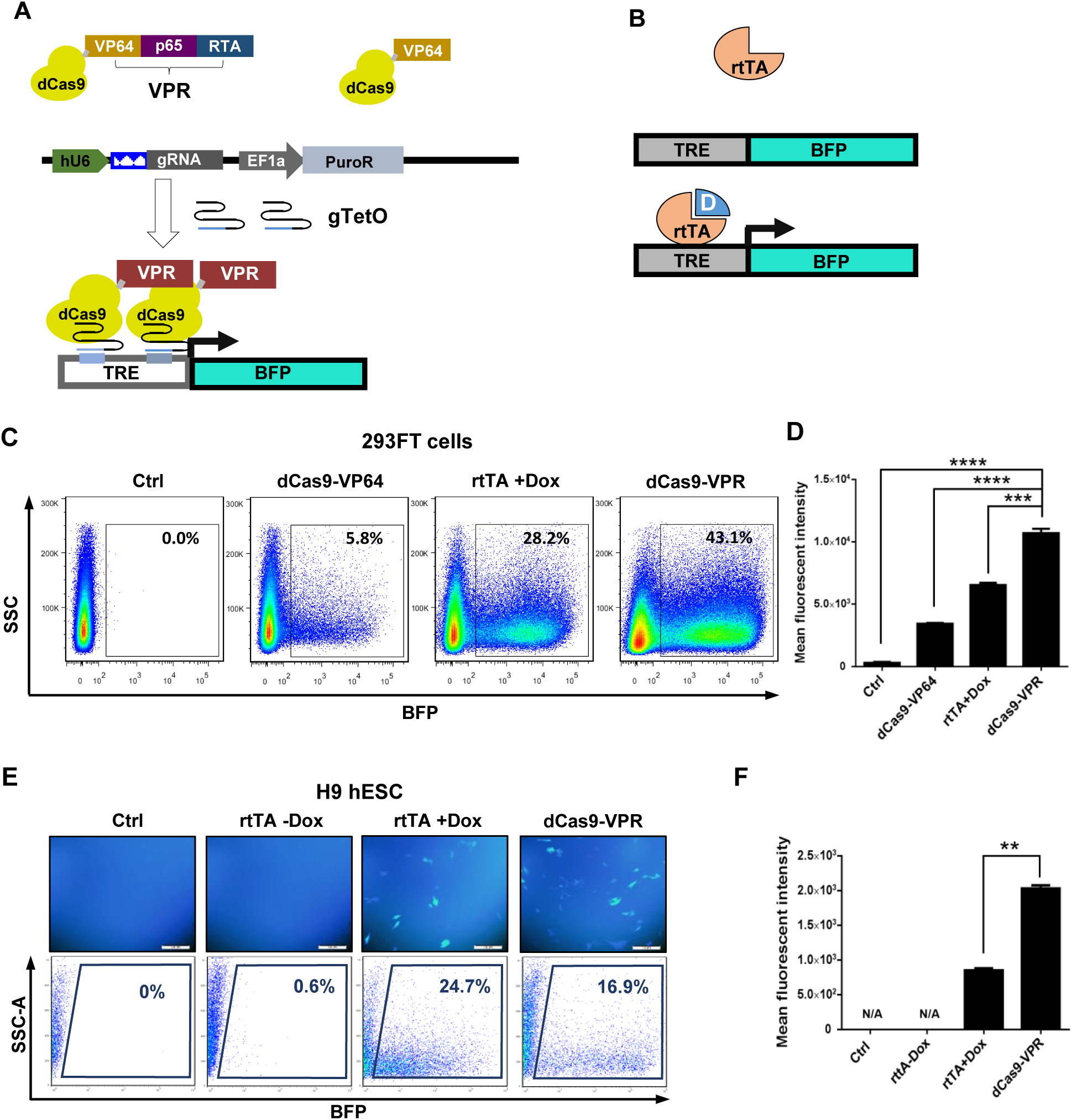
The dCas9-VPR system leads to robust transcription activation in human cell lines. (A) Schematic diagram of the gRNA guided dCas9-VPR gene activation system that consists of two parts: one plasmid contains dCas9-VPR driven by a CAG promoter; another plasmid contains gRNA targeting the promoter of the gene of interest driven by the human U6 promoter, in this case gTetO, and a PuroR selection cassette driven by an EF1α promoter. Upon co-transfection of the two plasmids, dCas9-VPR can activate the BFP transcription downstream of the TRE promoter. (B) Tet-On system: rtTA protein can bind to the TRE promoter and drive expression of the down-stream BFP gene in the presence of Dox. (C) 293FT cells were transfected with the reporter plasmid containing BFP driven by the TRE promoter. They were either co-transfected with dCas9-VPR or dCas9-VP64 and gTetO plasmids, or with the CAG-rtTA plasmid. Dox was added immediately after transfection. Cells were harvested 2 days after transfection and the fluorescence was analyzed using flow cytometry. (D) Bar graph quantification of mean fluorescent intensity analyzed using the FlowJo software v7.6.1. \*\*\**p* <0.001, \*\*\*\**p* <0.0001, *n*=3. (E) H9 hESCs were electroporated with either rtTA or dCas9-VPR+gTetO plasmids together with the TRE-BFP plasmid. Dox was added immediately after electroporation. Cells were harvested 3 days after electroporation and analyzed using flow cytometry. (F) Bar graph quantification of the mean fluorescent intensity analyzed using the FlowJo software v7.6.1. N/A, not applicable. \*\**p* <0.01, *n*=2.

We next tested the dCas9-VPR function in hESCs. DCas9-VPR, gTetO, and TRE-BFP plasmids were co-transfected into H9 hESCs. In another group, rtTA and TRE-BFP plasmids were co-transfected. FACS analysis showed that nearly 17% of cells in the dCas9-VPR group turned on BFP, while 24.7% of cells in the rtTA group were BFP positive after Dox induction, and only 0.6% of cells exhibited BFP fluorescence without Dox (Fig. 1E). Interestingly, the dCas9-VPR group showed the strongest mean fluorescence intensity (Fig. 1F). This is consistent with our result based on 293FT cells and proves that dCas9-VPR is a robust transcription activator, even compared with rtTA. We also tested the activation effect of dCas9-VPR in mouse embryonic stem cells (mESCs) and mouse embryonic fibroblasts (MEFs) and obtained similar results (Fig. S1A and B).

We then tested the efficiency of dCas9-VPR to activate normally silenced pluripotency genes in human cells. Two gRNAs targeting the −254 and −144 positions upstream of the transcription start site (TSS) of the pluripotency gene *NANOG* were selected (Fig. 2A). A GFP-2A-Puromycin resistant gene expression cassette was placed after the gRNA cassette both to monitor the transfection efficiency and for selection (Fig. 2A). *NANOG* cannot be activated by gNANOG alone or by dCas9-VPR together with the control gTetO. However, introducing gNANOG and dCas9-VPR together could elevate the *NANOG* transcript level by up to 150-fold in 293FT cells, indicating that it has a robust gene activation function (Fig. 2C).

**Figure 2.**
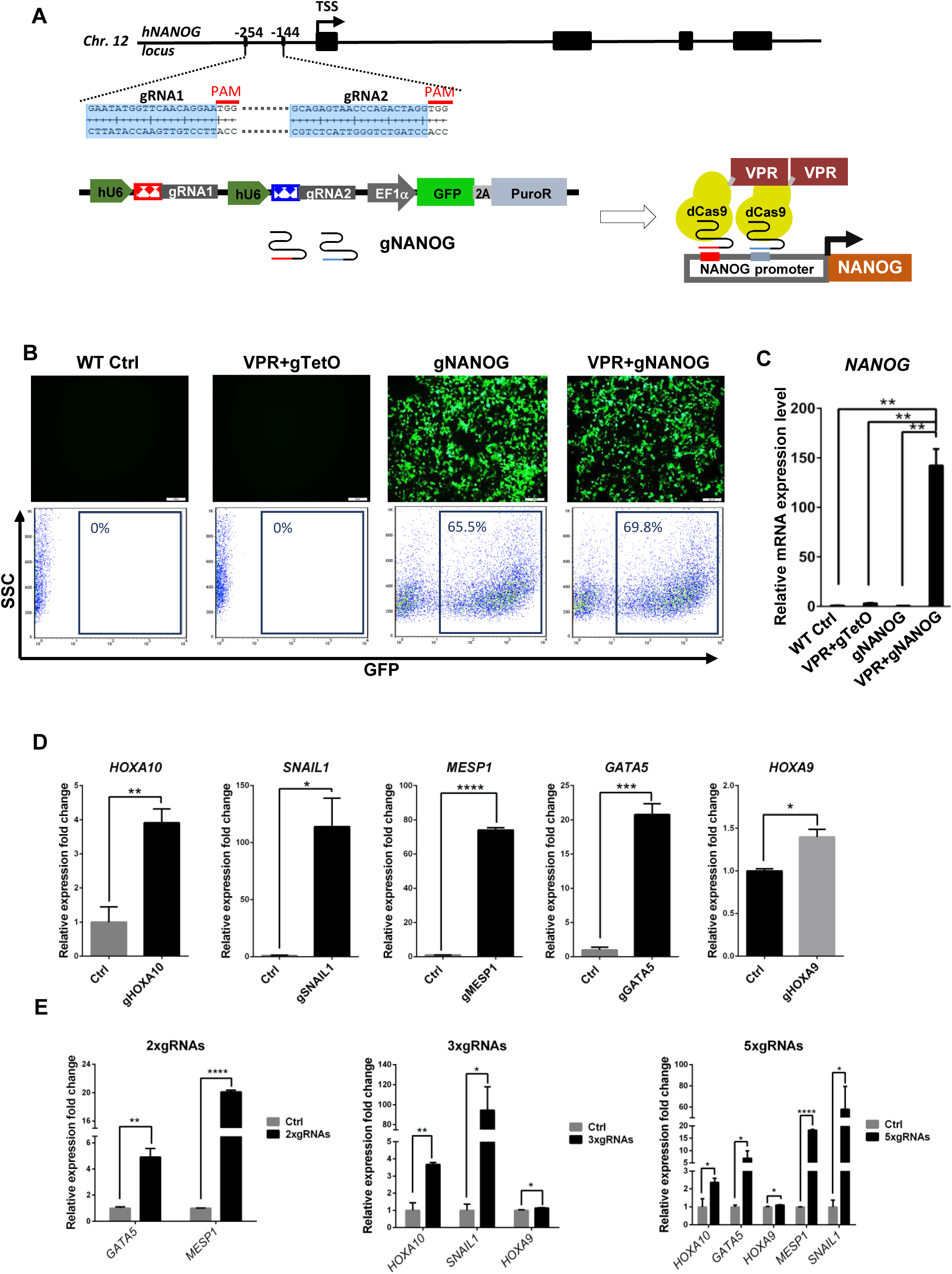
DCas9-VPR can be used to activate single or multiple genes in 293FT cells. (A) *NANOG* gRNA targeting sites were located at −254bp and −144bp upstream of the *NANOG* transcription starting site (TSS); protospacer-adjacent motif (PAM) sequences in red; black boxes indicate exons. (B) DCas9-VPR and gNANOG plasmids were co-transfected into 293FT cells. DCas9-VPR and gTetO plasmids were used as control. Top panels, fluorescence images of transfected cells; gNANOG plasmid transfected cells showed strong GFP fluorescence. Bottom panel, flow cytometry analysis of GFP^+^ cells in each group. (C) Q-PCR analysis of *NANOG* expression 2 days after transfection; the dCas9-VPR system showed nearly 150-fold up-regulation of *NANOG* mRNA. Relative gene expression values were normalized against *GAPDH*. Error bars represent SEM. \*\**p* <0.01, *n*=3. (D) Activation of endogenous genes by dCas9-VPR. DCas9-VPR was co-transfected with gRNA pairs targeting *HOXA10, SNAIL1, MESP1, GATA5* or *HOXA9*, respectively. Cells were harvested 2 days after transfection and subjected to Q-PCR analysis. All tested genes showed significant upregulation compared to the control group. All expression levels were normalized against *GAPDH*. Error bars represent SEM. \**p* < 0.05, \*\**p* < 0.01, \*\*\**p* < 0.001, \*\*\*\**p* < 0.0001, *n*=3. (E) Simultaneously activation of multiple endogenous genes in 293FT cells. DCas9-VPR was co-transfected with 2×gRNAs (gMESP1, gGATA5), 3×gRNAs (gHOXA10, gSNAIL1, gHOXA9) or 5×gRNAs (gHOXA10, gSNAIL1, gMESP1, gGATA5 and gHOXA9). Cells were harvested 2 days after transfection. Q-PCR analysis confirmed co-upregulation of multiple genes targeted by pooled gRNAs. All expression levels normalized against *GAPDH*. Error bars represent SEM. \**p* < 0.05, \*\**p* < 0.01, \*\*\*\**p* < 0.0001, *n*=3.

Next, we tested whether the dCas9-VPR system could simultaneously activate multiple genes in human cells, we designed 2 different gRNAs per gene promoter for *HOXA10, SNAIL1, MESP1 GATA5* and *HOXA9*. First we tested the activation efficiency of these gRNAs towards their target genes when transfected separately in 293FT cells (Fig. 2D). Q-PCR analysis showed all of the five pairs of gRNAs can activate their target gene upon co-transfection with dCas9-VPR (Fig. 2D). We next pooled gRNA pairs of two genes (2×gRNAs: *MESP1*, *GATA5*), three genes (3×gRNAs: *HOXA10*, *SNAIL1*, *HOXA9*) or five genes (5×gRNAs: *HOXA10*, *SNAIL1*, *MESP1*, *GATA5* and *HOXA9*) to test the co-activation efficiency. Upon co-transfection with dCas9-VPR, different combination of gRNAs upregulated their target genes together (Fig. 2E), indicating that dCas9-VPR system could be a useful tool for multiplexed endogenous gene activation.

To validate the utility of the dCas9-VPR system in hESCs, we transfected H9 hESCs with either dCas9-VPR and gNANOG or with rtTA and *NANOG* coding DNA sequence (CDS) joined to H2B-mCherry through a 2A peptide driven by a TRE promoter. As shown in Fig. 3A, for the dCas9-VPR group, increased NANOG protein expression (in red) can be detected in colonies with GFP fluorescence. Upon Dox induction, stronger NANOG was also visible in Tet-On system transfected cells and co-localized with the H2B-mCherry (Fig. 3A). Quantitative PCR (Q-PCR) and western blot confirmed the elevated NANOG level induced by either dCas9-VPR+gNANOG or *NANOG* CDS. The transcript level of another pluripotency marker gene, *OCT4*, was increased synergistically (Fig. 3B). Western blot analysis confirmed the upregulation of NANOG and OCT4 proteins in transiently transfected H9 cells (Fig. 3C). We generated a transgenic hESC line constitutively expressing dCas9-VPR and observed no cytotoxicity, decrease in pluripotency gene expression, or change in cell morphology for long-term cultures (Fig. 3D and E). This suggests that the dCas9-VPR system is suitable for gene activation studies in hPSCs.

**Figure 3.**
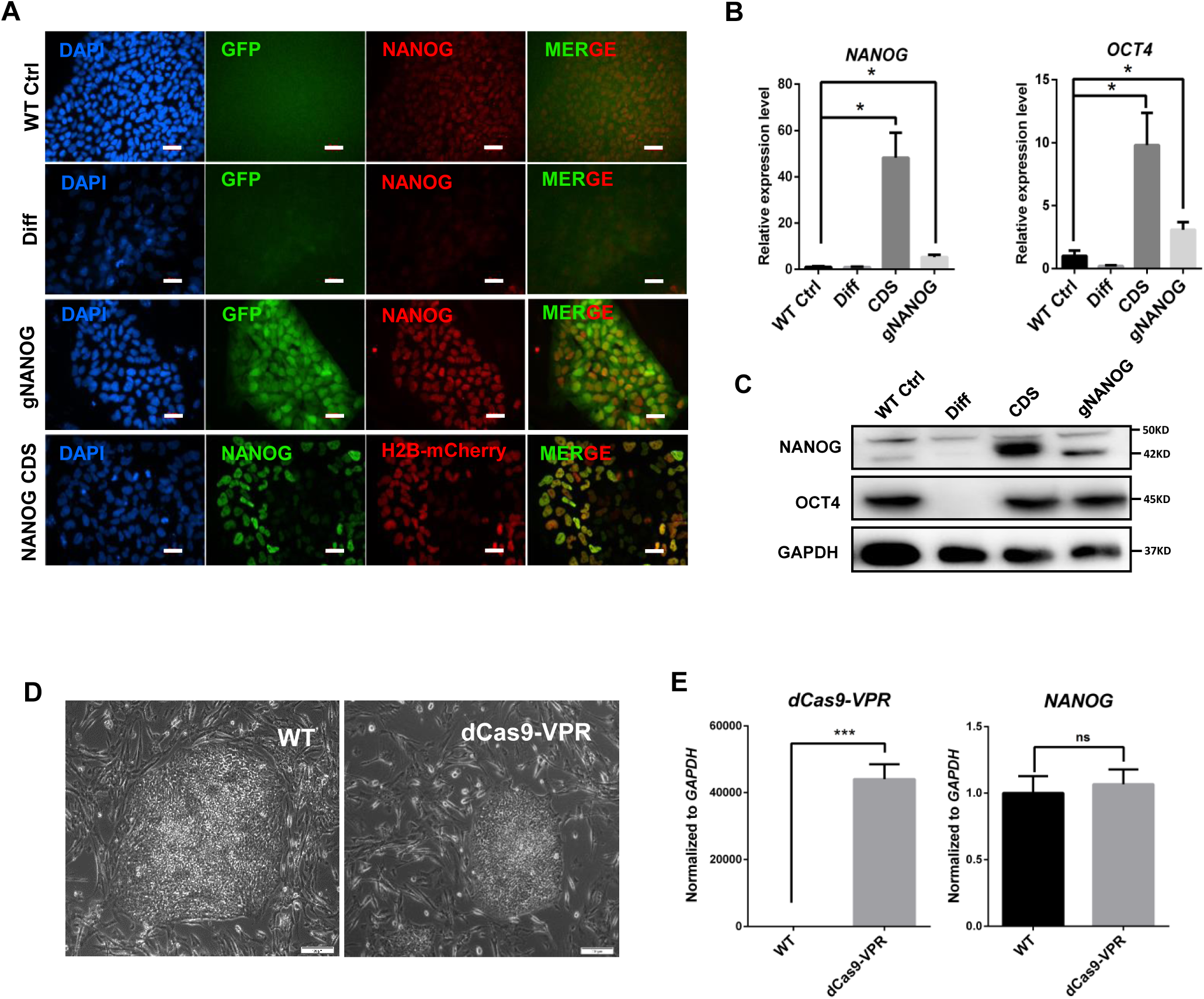
Activation of endogenous *NANOG* gene in hESCs by dCas9-VPR. (A) Immunostaining showing upregulation of NANOG protein by the dCas9-VPR system. Cells were fixed 5 days after transfection. WT Ctrl, untransfected H9 cells; Diff, differentiated H9 cells induced by 10 μM retinoic acid (RA); gNANOG, H9 cells co-transfected with dCas9-VPR and gNANOG plasmids; *NANOG* CDS, cells co-transfected with CAG-rtTA and TRE driving NANOG-2A-H2B-mCherry. Dox were added immediately after electroporation. All plasmids were based on the PiggyBac system and co-transfected with a plasmid containing HyperPB transposase driven by a CAG promoter. Scale bar, 20 μm. (B) Q-PCR analysis of *NANOG* and O*CT4* expression in H9 cells 5 days after transfection. All expression levels normalized against *GAPDH*. Error bars represent SEM. \**p* < 0.05, *n*=3. (C) Western blot analysis of NANOG and OCT4 protein expression in H9 hESCs. Cells were harvested 5 days after transfection without selection. (D) DCas9-VPR constitutive expressing H9 cells showed similar clone morphology after long-term culture. Scale bar, 100 μm. (E) Q-PCR result showing dCas9-VPR constitutive expressing H9 cells and wild-type H9 cells expressed similar amount of *NANOG*. All expression levels normalized against *GAPDH*. Error bars represent SEM. ns. *p* >0.05, \*\*\**p* <0.001, *n*=3.

### Generation of an inducible idCas9-VPR hESC knock-in line

To achieve efficient, tunable, and reversible gene activation while avoiding compromising the genome integrity of hPSCs, we engineered an iVPR system by inserting the CAG promoter driving the rtTA expression cassette and the TRE promoter driving the dCas9-VPR cassette into the two alleles of the *AAVS1* locus on chromosome 19. H9 hESCs were co-transfected with two donor plasmids containing dCas9-VPR and M2rtTA, as well as a pair of Cas9 nickase plasmids with *AAVS1* targeting gRNAs to induce DNA double-strand break (DSB) and homology recombination (HR) (Fig. S2A). After puromycin and neomycin double selection for 2 weeks, we picked and expanded 17 clones. Upon addition of Dox, all the clones showed clear induction of dCas9-VPR protein expression (Fig. S2B). Genomic DNA PCR was performed to select correct targeted clones and rule out random insertions (Fig. S2C). Clone 2, 6 and 8 had targeted insertion at both *AAVS1* alleles and without any random insertion (Fig. S2C). They were chosen for further analysis. Southern blot confirmed that in all three clones, both alleles of *AAVS1* contained the correct insertion (Fig. 4A and B). Q-PCR analysis showed that in hESCs, without Dox treatment, little *dCas9-VPR* transcript could be detected, while after Dox addition, strong *dCas9-VPR* expression was induced (Fig. 4C). Karyotype analysis showed that all three clones had normal 46XX karyotype (Fig. S2D). IVPR clone 2 was chosen for further study. Without Dox, we could not detect any dCas9-VPR protein in iVPR cells. The dCas9-VPR protein appeared after 12 hours of Dox addition and reached a plateau at 24 hours (Fig. 4D). While 6 hours after Dox withdrawal, the dCas9-VPR protein decreased, by 12 hours, it decreased to a low level and could not be detected anymore after 24 hours (Fig. 4D). The induction of dCas9-VPR from the *AAVS1* locus was not affected by differentiation. We induced mesoderm differentiation by culturing cells in an RPMI medium supplemented with albumin, ascorbic acid, transferrin, selenite, BMP4 (5ng/ml) and CHIR99021 (2 μM) as described by Burridge et al. (Burridge et al. 2015). Q-PCR analysis showed that after 3 days of differentiation, pluripotency marker genes *OCT4* and *SOX2* were significantly downregulated, while *dCas9-VPR* was highly expressed as long as Dox was present, regardless whether cells were in hESC culture medium E8 or in the differentiation medium (Fig. 4E). Genes related to mesoderm differentiation and epithelial to mesenchymal transition, such as *SNAIL*, were strongly upregulated by BMP4 and CHIR99021, confirming that hESCs had taken a mesoderm fate (Fig. 4E). These results suggest that the iVPR hESC line can be used for efficient and reversible gene activation.

**Figure 4.**
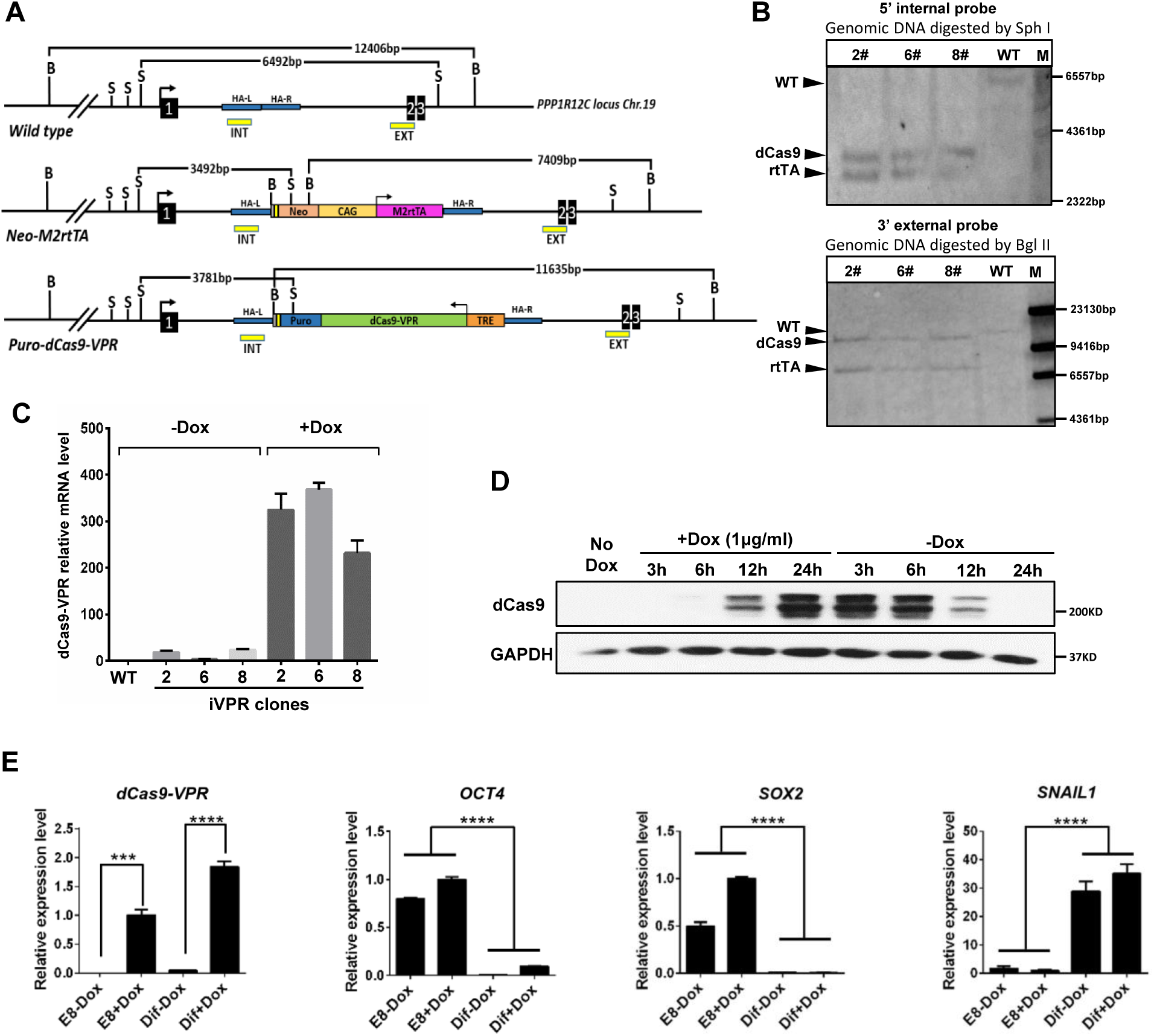
Generation of the iVPR hESC line. (A) Schematic view of wild type, targeted *AAVS1* locus, and positions of southern blot probes. B (Bgl II site), S (Sph I site), EXT (external probe), INT (internal probe). The sizes of the expected bands are indicated at the top. Blue lines indicate homology to the *PPP1R12C* intron. HA-L and HA-R, left and right homology arms. (B) Southern blot confirmed the correct targeted *AAVS1* locus in the iVPR clone 2#, 6#, 8#. M, marker. (C) Q-PCR analysis of dCas9-VPR transcript levels with or without Dox treatment. Expression levels were normalized against *GAPDH*. Error bar represents SEM. (D) Western blot of *dCas9-VPR* protein level upon Dox addition and after Dox withdrawal in idCas9-VPR clone 2. The time points are indicated at the top. (E) Q-PCR showing that the induction of *dCas9-VPR* was not affected by differentiation. Cells were induced to undergo mesoderm differentiation for 3 days in the presence or absence of Dox. Gene expression levels were all normalized against *GAPDH*. Error bar indicates SEM. \*\*\**p* < 0.001, \*\*\*\**p* < 0.0001, *n*=3.

### Upregulation of NANOG by dCas9-VPR promoted naïve state of pluripotency

The iVPR system provided a unique platform to investigate gene functions through activation from the endogenous locus. *NANOG* is a key regulator of pluripotency. We generated iNANOG hESCs by transfecting the PiggyBac based gNANOG plasmid described earlier into iVPR clone 2, 6, and 8, followed by FACS selection of GFP^+^ cells. Q-PCR analysis showed that after 2 days of Dox treatment, only iNANOG cells showed a significant increase (about 18 folds) in the *NANOG* mRNA level, while iVPR cells, with or without Dox, or iNANOG cells without Dox did not show any change in *NANOG* expression, indicating that the iNANOG system is tightly regulated (Fig. 5A). We also tested the time window of *NANOG* down-regulation after Dox withdrawal. *NANOG* mRNA was unchanged during the first 12 hours and decreased after 24 hours. It approached the background level after 48 hours (Fig. 5B). We next examined the change in NANOG protein level after Dox addition and withdrawal. Western blot revealed that dCas9-VPR protein became detectable 12 hours after Dox induction and reached a significant level after 24 hours (Fig. 5C, dCas9, long exposure; LE). Accordingly, NANOG protein showed an obvious increase after 24 hours and maintained at high level as long as dCas9-VPR was present (Fig. 5C, NANOG, LE). On the other hand, 6 hours after Dox removal, the dCas9-VPR protein decreased significantly (Fig. 5D, dCas9, short exposure; SE). The decline of the dCas9-VPR protein was most apparent during the first 24 hours. After 4 days without Dox, dCas9-VPR protein became almost undetectable (Fig. 5D). Similarly, the NANOG protein level dropped to the background level after 4 days of Dox withdrawal (Fig. 5D). Q-PCR analysis showed that after Dox induction, iNANOG significantly upregulated naïve state related genes such as *OCT4*, *PRDM14*, *GDF3*, and *LEFTYB*, while the early differentiation genes such as *AFP* was significantly downregulated (Fig. 5E). *XIST*, a long non-coding RNA involved in X chromosome inactivation were also downregulated after *NANOG* induction (Fig. 5F). The expression of SSEA3, a more rigorous pluripotency cell surface marker, was increased and became more homogeneous after *NANOG* elevation (Fig. 5G and Fig. S3). In addition to elevated expression of pluripotency genes, iNANOG cells also showed enhanced survival and proliferation abilities. Clonogenicity assay showed that after Dox induction, twice as many clones formed from dissociated iNANOG single cells (Fig. 5H and I). Finally, we tested whether *NANOG* upregulation by iVPR may facilitate hESCs to enter the naïve state of pluripotency. IVPR cells and iNANOG cells were cultured in 2iL medium which supplemented with ERK inhibitor PD0325901, GSK3 inhibitor CHIR99021, human LIF, and bFGF proteins with or without Dox addition (Silva et al. 2009; Takashima et al. 2014). Upon changing to the 2iL medium, hESCs colonies changed into a domed-shaped morphology and became more compact (Fig. 5J and K). INANOG cells without induction can only survive for no more than three passages in the 2iL medium (Fig. 5J). Interestingly, Dox induced iNANOG cells can grow in the 2iL medium for longer than 9 passages with single cell dissociation and a 1:15 passage ratio (Fig. 5J and K). In contrast to iNANOG cells, Dox treated iVPR cells could not survive in 2iL conditions (Fig. 5J). Thus, upregulation of *NANOG* from its endogenous locus significantly improved single cell clonogenicity and permitted hESCs to grow in a naïve state culture environment.

**Figure 5.**
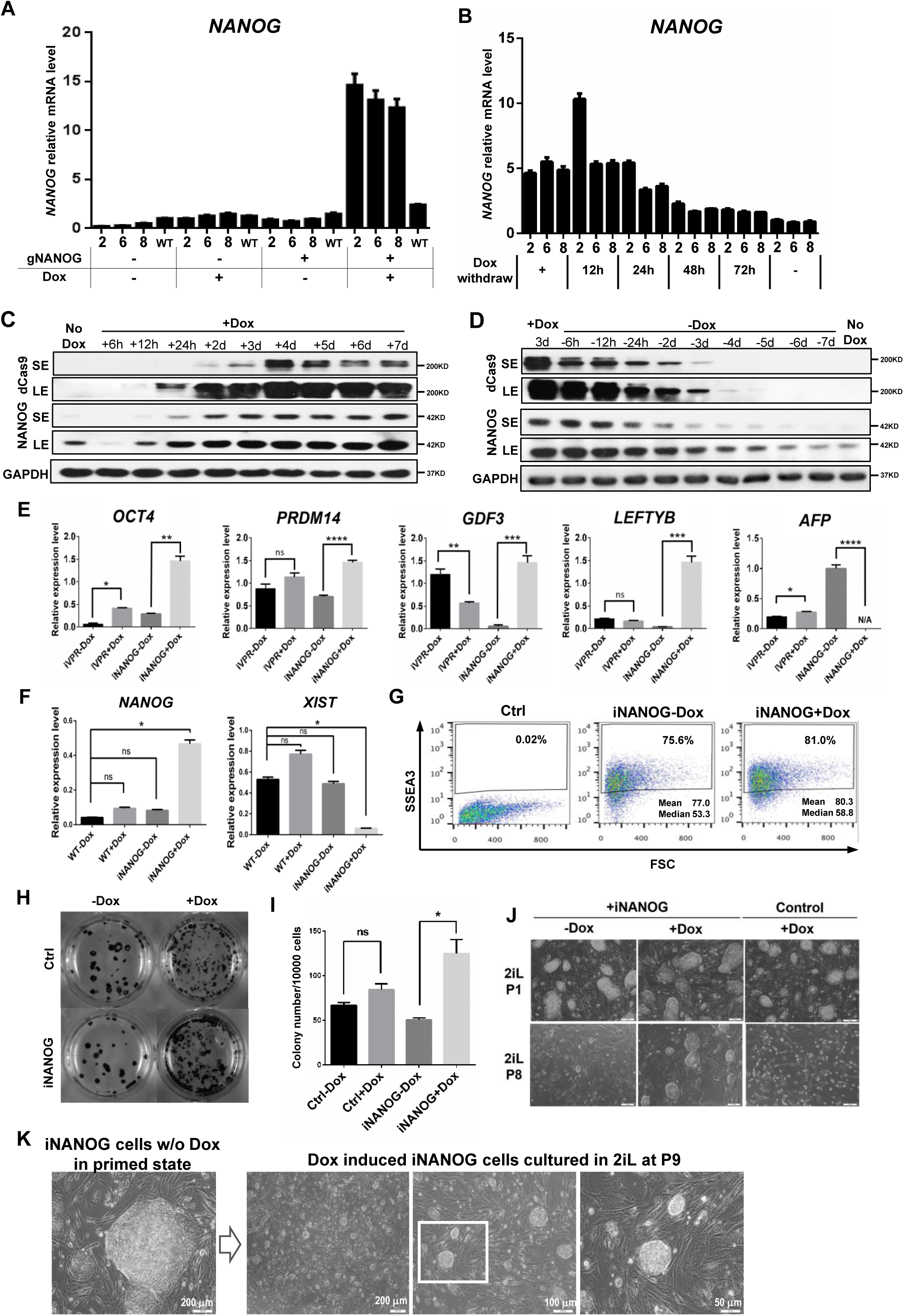
Upregulation of *NANOG* by dCas9-VPR promoted clonogenicity and the naïve state of pluripotency. (A) Q-PCR analysis of *NANOG* upregulation in iNANOG cells. IVPR clones 2, 6, and 8 were electroporated with gRNA expression plasmid targeting the *NANOG* promoter, as shown in Fig. 2A. GFP positive cells were purified by FACS and maintained as iNANOG cells. They were treated with or without Dox (1 μg/ml) for 2 days. *NANOG* expression level was normalized against *GAPDH*. Error bar represents SEM. *NANOG* expression level significantly elevated only in iNANOG+Dox group. n=3. (B) Q-PCR analysis of *NANOG* down-regulation in iNANOG cells. Dox was added for 2 days, then removed. Cells were harvested at different time points, as indicated. *NANOG* expression was normalized against *GAPDH*. Error bar represents SEM. *NANOG* expression level significantly dropped during Dox withdraw. *n*=3. (C) Western blot showing increased NANOG protein expression in iNANOG cells at different time points after Dox treatment. SE, short exposure; LE, long exposure; d, day; h, hour. (D) Western blot showing NANOG protein expression decrease in iNANOG cells at different time points after Dox withdrawal. SE, short exposure; LE, long exposure; d, day; h, hour. (E) Q-PCR analysis showing upregulation of pluripotency gene *OCT4, PRDM14, GDF3*, and *LEFTYB*, and down-regulation of differentiation gene *AFP.* Expression level all normalized against *GAPDH*. Error bar represents SEM. N/A, not applicable. ns. *p* >0.05, \**p* < 0.05, \*\**p* < 0.01, \*\*\**p* < 0.001, \*\*\*\**p* < 0.0001, *n*=3. (F) Q-PCR analysis showing downregulation of *XIST* after *NANOG* induction. Expression level normalized against *GAPDH*. Error bar represents SEM. ns. *p* >0.05, \**p* < 0.05, *n*=3. (G) Flow cytometry analysis showing increased SSEA3 expression after *NANOG* induction. Data analyzed using the FlowJo software v7.6.1. (H) Clonogenicity assay of iNANOG cells. Alkaline phosphatase assay (dark blue) was used to visualize undifferentiated colonies. (I) Bar graph quantification of the clonogenicity assay. ns. *p* >0.05, \**p* < 0.05, *n*=3. (J) Morphology of iNANOG cells cultured in the 2iL medium. *NANOG* overexpression (iNANOG + Dox) promoted long-term cell growth in the 2iL medium. Representative images of passages 1 and 8 (P1 and P8) are shown. Scale bar, 100 μm. (K) Morphology of primed state iNANOG cells (without Dox) and Dox induced iNANOG cells (passage 9, P9) in the 2iL medium.

### Upregulation of *NANOG* enabled hESCs to integrate with mouse ICM *in vitro*

Entering the pluripotent ICM lineage is considered a more stringent test for naïve state ESCs (Gafni et al. 2013; Takashima et al. 2014). We next used *in vitro* human–mouse blastocyst chimera assay to assess the functionality of iNANOG cells (Fig 6A). To exclude the influence of Dox treatment only, wild type hESCs stably carrying gNANOG (WTSG) were used as the control. For this series of experiments, we also added Forskolin (a cAMP agonist) into the 2iL medium, since it had been shown to promote hPSCs to enter the naïve state (Hanna et al. 2010; Ware et al. 2014; Duggal et al. 2015). We refer to this medium as 2iL/FK. iNANOG cells showed further enhanced proliferation in the 2iL/FK medium and were able to form large, dome-shaped colonies (Fig. 6B), while cells without *NANOG* overexpression could only form small colonies (Fig. S4A). E3.5 blastocysts were collected from ICR mice for hESC injection. iNANOG cells and WTSG cells cultured with or without Dox, in either the E8 or 2iL/FK medium, were dissociated into single cells. 10–15 single cells were injected into the blastocoel cavity and cultured in a 1:1 mixed KSOM:2iL/FK medium for 24 hours (Fig. 6A). Because cells without *NANOG* overexpression only formed small colonies on feeder in the 2iL/FK medium, we could not obtain sufficient pure hESCs for blastocyst injection. Therefore, this group was omitted from this series of experiments. Since all cells used for injection contained GFP transgene expressed from the gNANOG plasmid, the location of human cells in the mouse blasocysts could be followed directly under the fluorescence microscope. 4–6 hours after injection, most blastocysts contained GFP positive human cells (Fig. 6B and C). After 24 hours of culture, many embryos still contained hESCs (Fig. 6B). We used time-lapse imaging to monitor the activity of hESCs in mouse blastocysts over time (Supplementary movie S1). Interestingly, endogenous *NANOG* overexpression strongly enhanced the survival of hESCs in mouse blastocysts. 12 hours after injection, 2iL/FK cultured Dox induced gNANOG cells could be found in approximately 82% of blastocysts, while E8 cultured Dox induced gNANOG cells were alive in 73% of blastocysts (Fig. 6C and S4B). In contrast, without Dox induction, E8 cultured iNANOG cells could only be seen in 49% of injected blastocysts (Fig. 6C and S4B). We next analyzed the locations of the transplanted hESCs. Injected embryos were fixed after 24 hours of culture, stained with CDX2 (a trophectoderm marker) and β-Catenin, and observed with a confocal microscope. Different integration patterns were shown: hPSCs integrated into the ICM region (ICM), in both the ICM and the trophectoderm (Multiple), only in the trophectoderm (TE), and disappeared (None) (Fig. 6D). We also performed live imaging to monitor the behavior of iNANOG cells in the mouse blastocyst. Interetingly, 2iL/FK cultured iNANOG cells tend to migrate with mouse inner cell mass cells as blastocyst hatching from the zona pellucida (Fig. 6E). *NANOG* overexpression significantly improved the percentage of cells remaining in blastocysts, and the 2iL/FK culture further increased the ICM integration proportion (Fig. 6F, G, and S4C). On average, two 2iL/FK or E8 cultured iNANOG cells could be found in the ICM region 24 hours after injection, while without *NANOG* overexpression, hardly any GFP cells were seen in the ICM (Fig. 6G). Thus, upregulation of *NANOG* from its endogenous locus greatly enhanced cell survival and their subsequent ICM integration in hPSC-mouse blastocyst chimeras.

**Figure 6.**
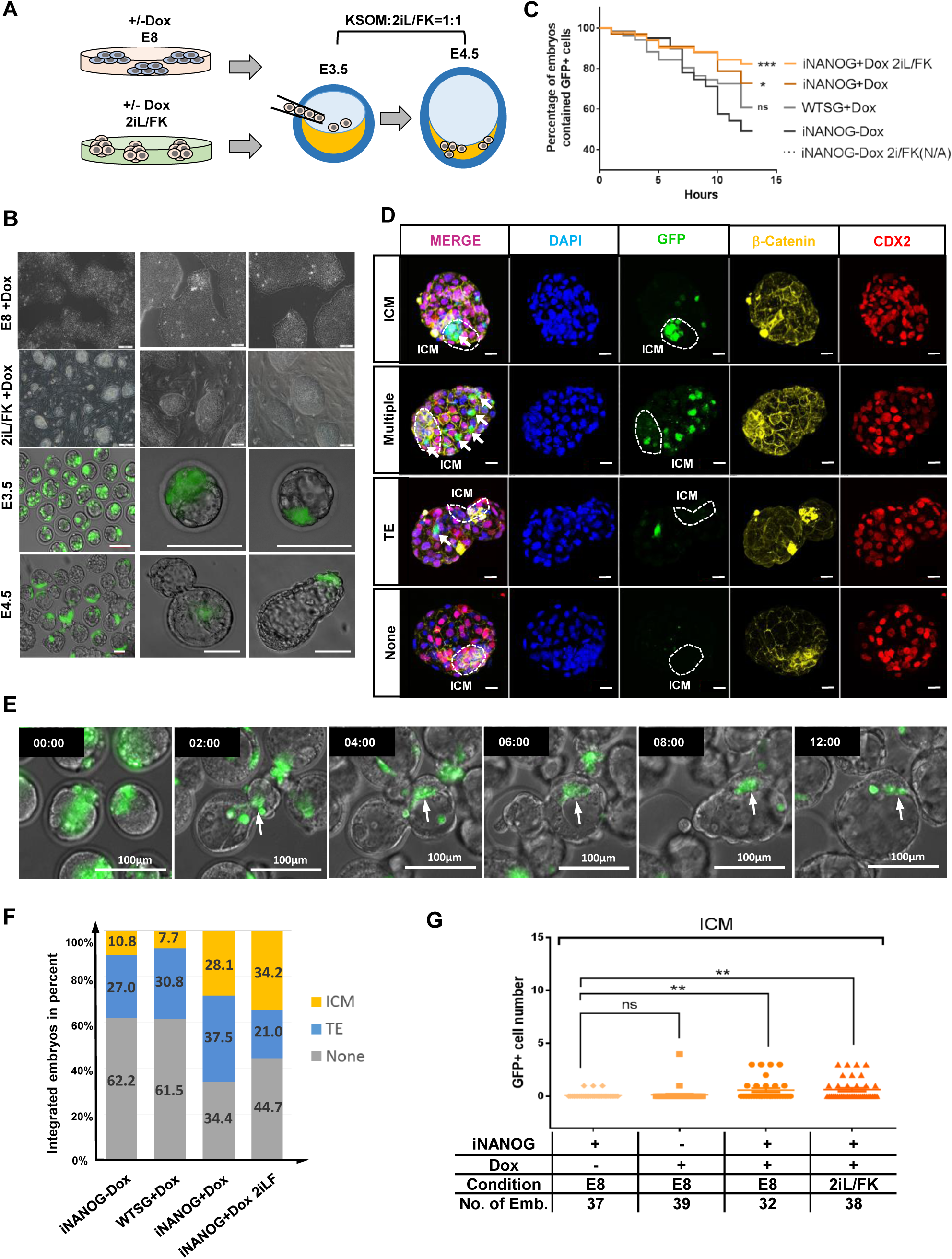
Upregulation of *NANOG* by idCas9-VPR promoted hESC survival and ICM integration in mouse blastocysts *in vitro*. (A) Cartoon showing *in vitro* hESC-mouse blastocyst chimera formation assay. iNANOG cells were cultured in E8 or 2il/FK medium with or without Dox, then injected into E3.5 mouse blastocysts and cultured to E4.5 in KSOM: 2il/FK = 1:1 medium *in vitro*. (B) Morphology of iNANOG cells in culture and chimeric embryos. Top 2 rows, cells cultured in E8 or 2iL/FK on feeders; scale bar, 100 μm. Bottom 2 rows, E3.5 and E4.5 mouse blastocysts with iNANOG cells (GFP); scale bar, 100 μm. (C) Survival curve of hESC in mouse blastocysts over time. WTSG, wild type hESCs expressing *NANOG* gRNA. The *p* value was calculated using the Log-rank (Mantel-Cox) test. ns. *p* >0.05, \**p* < 0.05, \*\*\**p* < 0.001. Detailed information is provided in Fig. S4B. (D) Confocal images of E4.5 chimeric embryos. β-Catenin, yellow; CDX2, red; DNA, blue; iNANOG cells, green. The ICM region is highlighted by a dashed circle. Scale bar, 20 μm. (E) Selected frame from time-lapse movie of iNANOG-mouse blastocyst chimera. Arrows indicating iNANOG cells moved with mouse inner cell mass cells during blastocyst hatching. Scale bar, 100 μm. (F) The proportion of blastocysts with hESC integration for E4.5 embryos. Blastocysts with GFP cells in both ICM and TE were counted as ICM. The percentage of embryos with ICM/TE/None integration was labeled in the colored bar. (G) Dot graph showing the number of iNANOG cells in the ICM region of E4.5 embryos. Cells, culture condition before injection, and number of embryos were as listed. Error bars represent SEM. ns. *p* >0.05, \*\**p* < 0.01.

## Discussion

In this study, we generated an inducible CRISPR-ON hESC line by targeting the *AAVS1* locus. Based on both our results and those of Chavez et al. (Chavez et al. 2015), dCas9-VPR appeared to be a stronger activator than VP64 to induce gene expression from both ectopic and endogenous promoters. It even led to a higher level of reporter gene activation compared with Tet-ON rtTA, where VP64 was fused with Tet protein directly bound to the TRE elements. This is likely due to the combined effects of VP64, NF-κB transcactivating subunit p65, and the viral transcription factor Rta, which together can recruit a multitude of endogenous factors to achieve dramatically enhanced transcriptional activation. Other dCas9 based transcription activators have been generated. For example, Balboa et al. found increased activation ability with more VP16 fusing together. Using the longest version of dCas9-VP192 combined with inducible systems, they sucessfully facilitated human cell reprogramming and differentiation (Balboa et al. 2015). Konermann et al. engineered a structure-guided CRISPR synergistic activation mediator system (SAM), where they engineered gRNA2.0 by replacing the tetraloop and stem loop 2 of the original gRNA with a minimal hairpin aptamer that specifically binds to MS2 bacteriophage coat proteins (Konermann et al. 2014). By co-expression of dCas9-VP64, gRNA2.0, and MS2 fused with p65 and the activation domain of the human heat-shock factor 1 (HSF1), highly effective gene activation can be achieved (Konermann et al. 2014). Tanenbaum et al. constructed a SunTag system: dCas9 was joined with 10 copies of GCN4 peptide (SunTag), while VP64 was fused with scFv-GCN4 (the single-chain variable fragment (scFv) antibody of GCN4) (Tanenbaum et al. 2014). When co-expressed in the cell, SunTag was bound by scFv-GCN4, and multiple copies of VP64 resulted activation of the target gene (Tanenbaum et al. 2014). Compared with the systems discussed above, which required introducing tandem repeat large cassette or the co-expression of two components in addition to the gRNA, dCas9-VPR is a simple and effective option.

In our study, we chose to insert the iVPR system into the *AAVS1* locus, since it has been used as a ‘safe habor’ for transgene insertion in human stem cell systems (Dekelver et al. 2010). For example, Genga et al. constructed a GFP labled H1 hESC line by knocking-in a CAG-GFP into the *AAVS1* locus. Besides, an inducible dCas9-KRAB gene inhibition system was also introduced into the GFP-H1 cells. By infecting sgRNA targeting the exdogenous CAG promoter, they successfully realized CRISPR based inhibition of exdogenous gene in hESCs (Genga et al. 2015). González et al. inserted the Dox inducible Cas9 system into the 2 alleles of the *AAVS1* locus of HUES8 hESCs (González et al. 2014; Zhu et al. 2014; Zhu et al. 2015). The resulting iCRISPR hESC line enabled selection-free gene knock-out and the generation of lineage-specific knock-in reporters. This demonstrated that when Cas9 was expressed in a controllable manner from a suitable locus, the resulting cell line can be a powerful platform for genome editing in normally hard to transfect human stem cells (Zhu et al. 2015). Similarly, using the iVPR line, we found that the efficiency to generate an iNANOG line was much improved. Upon Dox addition and withdrawal, *NANOG* transcripts and proteins could be up- and down-regulated in a highly repeatable manner, which greatly facilitated downstream experiments. Recently, Ordovás et al. reported *AAVS1*-locus mediated transgene inhibition in hESCs, and that inhibition may due to different cassettes inserted into the locus (Ordovás et al. 2015). We tested the iVPR expression in both undifferentiated hESCs and after induction of mesoderm differentiation. The level of *dCas9-VPR* transcripts was even higher upon Dox treatment after mesoderm induction (Fig. 4E). The iVPR and iNANOG cells have been maintained for more than 6 months, and we did not observe any reduction in the level of *dCas9-VPR* or *NANOG* induced by Dox. Thus, results us and other groups suggested that, in most cases, *AAVS1* locus integration is a reliable approach to generate transgenic hPSCs.

*NANOG* is a master transcription factor for pluripotency in both human and mouse ESCs (Mitsui et al. 2003; Boyer et al. 2005; Chambers et al. 2007). During somatic cell reprogramming to pluripotent stem cells, ectopic expression of *NANOG* helped to speed up reprogramming and restrict partially reprogrammed cells to the ground state (Hanna et al. 2009; Silva et al. 2009). Different from mESCs, conventional cultured hESCs are in a primed state, similar to the epiblast stem cells in mice (Brons et al. 2007; Tesar et al. 2007). Recently, multiple groups reported methods to obtain naïve state hPSCs that resemble ground-state mESCs (Gafni et al. 2013; Duggal et al. 2015; Takashima et al. 2014; Theunissen et al. 2014). Takashima et al. showed that ectopic expression of *NANOG* and *KLF2* could reset the self-renewal requirements of hPSCs so that they can be grown in a medium containing ERK1/2 inhibitor PD0325901 and GSK3 inhibitor CHIR99021, and adopt a domed-shaped morphology similar to that of mESCs (Takashima et al. 2014). Here we increased the expression of endogenous *NANOG* by targeting a strong transcription activator, dCas9-VPR, to its promoter. As expected, we observed upregulation of naïve state genes such as *GDF3*, *PRDM14*, and *LEFTYB* and downregulation of early differentiation gene *AFP*. Interestingly, these iNANOG cells showed a significantly improved survival ability and clonogenicity when cultured in the primed state, and they could grow in 2i plus LIF conditions for more than nine passages. The improved survival and self-renewal of iNANOG cells was not due to the effect of Dox treatment as described by Chang et al.(Chang et al. 2014), because Dox treated iNANOG cells showed significantly higher clonogenicity over Dox treated iVPR cells (Fig. 5H and I). The enhanced survival ability seemed to have a significant influence on whether hPSCs can integrate with the ICM of mouse blastocysts during *in vitro* culture. We found that even when iNANOG cells were in the primed state, after injection into mouse blastocysts, more cells remained inside the blastocysts and some of the cells were able to integrate with mouse ICM cells (Fig. 6F and G). Culturing iNANOG cells in 2iL/FK naïve state medium (Duggal et al. 2015) further improved the ICM integration rate (Fig. 6F and G). INANOG cells displayed highly dynamic interactions with mouse ICM cells, as observed in time-lapse movies (Supplementary movie S1). They migrated with mouse ICM cells as blastocysts hatched from zona pellucida. However, despite enhanced survival ability of iNANOG cells, many injected cells died over time. After 24 hours, more than 30% of injected blastocysts lost all iNANOG cells and more than 60% of blastocysts lost the injected hESCs if *NANOG* was not overexpressed (Fig. 6F). This was partially caused by poor survival of hESCs in the IVC1 and −2 media designed to culture peri-implantation mouse and human embryos (Bedzhov and Zernicka-Goetz 2014; Deglincerti et al. 2016; Shahbazi et al. 2016) (Fig. S4D). Thus, to achieve better naïve hPSC and mouse ICM integration, a culture medium suitable for both mouse blastocysts and hPSCs may be needed. The effect of *NANOG* overexpression on cell survival and self-renewal is also in accordance with the observation that chromosome 12, where the *NANOG* gene is located, is the most frequently gained chromosome in culture adapted hPSCs (Baker et al. 2007) and during hiPSC generation (Taapken et al. 2011). Moreover, *NANOG* was reported to be upregulated by a number of factors such as *STAT3*, Hedgehog signaling, hypoxia, etc., in human cancers, and repression or ablation of *NANOG* inhibited tumor initiation (Gong et al. 2015). Thus, iNANOG hESCs, where the endogenous *NANOG* can be activated by dCas9-VPR in a controllable manner, may also be a good system to study the process of hPSC adaptation and cancerous transformation.

In summary, the iVPR hESC line generated and characterized in this study offered a convenient, stable, and highly controllable platform for gene activation studies. It can also be used to investigate the function of regulatory elements in the genome such as super enhancers as well as for genome wide screens using established human gRNA libraries.

## Methods

### HESC culture

H9 hESCs (WiCell Institute) were maintained on inactivated mouse embryonic fibroblast (MEF) cells in standard hESC medium consisting of KO-DMEM (Invitrogen) supplemented with 1×Nonessential Amino Acids (NEAA) (Invitrogen), 0.1 mM 2-mercaptoethanol (Sigma-Aldrich), 1 mM GlutaMAX (Invitrogen), 20% Knock-out serum-replacement (KOSR) (Invitrogen) and 4 ng/ml bFGF (Peprotech). Cells were cultured at 37 ºC in a humidified atmosphere with 5% CO_2_ in air. They were passaged with 1 mg/ml collagenase IV (Invitrogen) and seeded onto MEFs. For feeder-free culture, hESCs were grown for more than three passages on Matrigel (growth factor reduced, BD Biosciences) in the absence of feeders in E8 medium (Invitrogen)

### Plasmid construction

DCas9-VPR was constructed by fusing the nuclease deficient Cas9 (dCas9) with transcription activator VP64, p65, and Rta in tandem as described by Chavez et al. (Maeder et al. 2013). For constitutive expression, dCas9-VPR was placed behind a CAG promoter in a PiggyBac vector also containing a PGK promoter driving a hygromycin resistance gene. For inducible expression from the *AAVS1* locus, dCas9-VPR was placed behind a TRE promoter in the *AAVS1* homologous recombineering donor plasmid, as shown in Fig. S2A. DCas9-VP64 was constructed by fusing dCas9 with VP64. Tet-On system was obtained from Clontech (http://www.clontech.com). PiggyBac plasmids were generous gift from the Sanger institute, Cambridge, UK (http://www.sanger.ac.uk). The multiple *NANOG* gRNA expression plasmid was constructed by SynGene (http://syngen.tech) as depicted in Fig. 2A.

### Naïve state culture condition for hPSCs

For naïve state conversion, cells cultured in standard hESC medium on MEFs were dissociated to single cells using 0.05% trypsin/EDTA solution (Invitrogen), replated on MEFs, and cultured overnight in standard hESC medium supplement with 10 μM Rho Kinase (ROCK)-inhibitor Y-27632 (Calbiochem). The next day, the standard medium was changed to the 2iL or 2iL/FK (for injection) medium, which consisted of KO-DMEM (Invitrogen), 20+ KOSR, 1×NEAA, 0.1 mM 2-mercaptoethanol, 1 mM GlutaMAX, 12 ng/ml bFGF, 10 ng/ml human recombinant LIF (Peprotech), 1 μM ERK1/2 inhibitor PD0325901 (Peprotech), 3 μM GSK3 inhibitor CHIR99021 (Peprotech), 10 μM Forskolin (Peprotech), and 50 μg/ml ascorbic acid (Sigma). HESCs changed to a dome-shaped morphology within 4–6 days after culturing in the 2iL or 2iL/FK medium and were passaged every 4 days as single cells using 0.05% trypsin/EDTA.

### Cardiac mesoderm differentiation from hESCs

For cardiac mesoderm differentiation, hESCs maintained on Matrigel (growth factor reduced, BD Biosciences) in E8 were dissociated into single cells with Accutase (Invitrogen), then seeded onto Matrigel-coated tissue culture dishes at a density of 5×10^4^ cells/cm^2^ and cultured in E8 for 3 days. Then the medium was switched to the RPMI1640 medium supplemented with Albumin, Ascorbic acid, transferrin, selenite, 5 ng/ml BMP4 (R&D Systems), and CHIR99021 to induce cardiac mesoderm formation.

### Quantitative PCR

Total RNA was extracted with TRIZOL (Invitrogen). 1 μg RNA of each sample was used for reverse transcription with Superscript III (Invitrogen). Q-PCR reactions were performed using GoTaq qPCR Master Mix (Promega) in a CFX96 Real-Time System (Bio-Rad). The relative expression level of each gene was normalized against the Ct (Critical Threshold) value of the house-keeping gene *GAPDH* using the Bio-Rad CFX Manager program. Primer sequences are listed in table S2.

### Antibodies, immunostaining, western blot, and FACS analysis

For immunostaining, cells were fixed in 4% paraformaldehyde (PFA) in PBS, permeabilized in 0.5% Triton X-100 (Sigma), blocked in 5% normal goat serum (Origene) and incubated with primary antibodies against NANOG (1:200), SSEA3 (1:200) in 4 °C overnight and detected by DyLight 488- or 549-conjugated secondary antibodies (Thermo). Nuclei were stained with DAPI (Sigma). A Nikon Ti-U fluorescence microscope was used for image acquisition. For western blot, cells were lysed in a RIPA buffer (Applygen, http://applygen.com.cn) with Protease Inhibitor Cocktail (Roche). Total proteins were separated on a 12% SDS/PAGE gel, transferred to nitrocellulose membrane (Whatman). The membrane was blocked with 5% non-fat dry milk in TBST and then incubated with primary antibodies against Cas9 (Genetex, 1:1000), GAPDH (CWBio, 1:1000), OCT4 (Santa Cruz, 1:1000) and NANOG (Cell Signaling Technology, 1:1000). After washing, the membrane was incubated with anti-mouse or anti-rabbit peroxidase-conjugated secondary antibodies (ZSGB-Bio http://www.zsbio.com/). Bands recognized by antibodies were revealed by ECL reagent (Pierce). For FACS analysis, cells were first dissociated with 0.05% Trypsin in 0.2% EDTA and PBS. FACS was performed on a Fortessa flow cytometer (Becton Dickinson).

### Mouse blastocyst injection and in vitro culture

Mouse morula were collected from ICR females 2.5 days post-coitus and cultured in KSOM medium (95mM NaCl, 2.5mM KCl, 0.35mM KH_2_PO_4_, 0.2mM MgSO_4_·7H_2_O, 0.2mM glucose, 10mM sodium lactate, 25mM NaHCO_3_, 0.2mM sodium pyruvate, 1.71mM CaCl·2H_2_O, 0.01mM EDTA, 1mM L-glutamine, 0.1mM EAA, 0.1mM NEAA, 4mg/ml BSA) at 37℃，5%CO_2_ for 24 hours to get blastocysts. HESCs were briefly treated with Accutase for single cell and injected (~10-15 cells for each embryo) on a Nikon microscope fitted with piezo-driven Eppendorf NK2 micromanipulator, CellTram air and CellTram Vario. After injection, embryos containing hESCs were cultured in medium supplemented with naïve culture medium：KSOM (1:1) (Chen et al. 2015) in 37 °C, 5% CO_2_ incubator. After injected embryos reformed blastocoel, the chimera embryos were live cell imaged using Leica microscope fitted with a live cell imaging system and fixed after 24-36 hours post-injection for staining and confocal imaging. For embryo immunostaining, zona pellucida-free injected embryos were fixed with 3.5% paraformaldehyde, permeabilized in 0.5% Triton X-100 (Sigma) and blocked with 5% BSA and then incubated with primary antibodies against CDX2 (BioGenex), β-Catenin (1:50, Abcam) and detected by DyLight 549- or 633- conjugated secondary antibodies(Thermo). Nuclei were stained with DAPI (Sigma). A Nikon-A1 fluorescence microscope was used for image acquisition.

### Statistical analysis

Data are presented as mean ± standard error of the mean (SEM). Statistical significance was determined by Student’s *t*-test (two-tail) for two groups or one-way Analysis of Variance (ANOVA) for multiple groups using Graphpad software. *p* < 0.05 was considered significant.

## Acknowledgments

This work was supported by the National Basic Research Program of China, 973 program grant 2012CB966701, the National Natural Science Foundation of China (NSFC), grant 31171381 (to J. N.), and core facilities of the Tsinghua–Peking University Center for Life Sciences. TNLIST Interdisciplinary research foundation grant 042003171 (to Z.X. and N.J.). We thank Dr Danwei Huangfu for the *AAVS1* homologous recombineering donor plasmids and Dr Xiaohua Shen lab for assistance in southern blot.

## Author contributions

J.G.: concept and design, collection and/or assembly of data, data analysis and interpretation, manuscript writing; D.M., R.H., M.Y., J.M.: collection and/or assembly of data; K.K. provided essential reagents, technical and scientific advice to the experiments and manuscript; Z.X.: concept and design; J.N.: concept and design, manuscript writing, and final approval of the manuscript. The authors declare no conflicts of interest.

## Compliance with ethical guidelines

Jianying Guo, Dacheng Ma, Rujin Huang, Jia Ming, Min Ye, Kehkooi Kee, Zhen Xie, and Jie Na declare that they have no conflict of interest.

This article does not contain any studies with human subjects performed by the any of the authors. All institutional and national guidelines for the care and use of laboratory animals were followed.

